# Hadza *Prevotella* Require Diet-derived Microbiota Accessible Carbohydrates to Persist in Mice

**DOI:** 10.1101/2023.03.08.531063

**Authors:** Rebecca H Gellman, Matthew R Olm, Nicolas Terrapon, Fatima Enam, Steven K Higginbottom, Justin L Sonnenburg, Erica D Sonnenburg

## Abstract

Industrialization has transformed the gut microbiota, reducing the prevalence of *Prevotella* relative to *Bacteroides*. Here, we isolate *Bacteroides* and *Prevotella* strains from the microbiota of Hadza hunter-gatherers of Tanzania, a population with high levels of *Prevotella*. We demonstrate that plant-derived microbiota-accessible carbohydrates (MACs) are required for persistence of *Prevotella copri* but not *Bacteroides thetaiotaomicron in vivo*. Differences in carbohydrate metabolism gene content, expression, and *in vitro* growth reveal that Hadza *Prevotella* strains specialize in degrading plant carbohydrates, while Hadza *Bacteroides* isolates use both plant and host-derived carbohydrates, a difference mirrored in *Bacteroides* from non-Hadza populations. When competing directly, *P. copri* requires plant-derived MACs to maintain colonization in the presence of *B. thetaiotaomicron*, as a no MAC diet eliminates *P. copri* colonization. *Prevotella’s* reliance on plant-derived MACs and *Bacteroides’* ability to use host mucus carbohydrates could explain the reduced prevalence of *Prevotella* in populations consuming a low-MAC, industrialized diet.

**Statement on work with indigenous communities:** In order to acquire scientific knowledge that accurately represents all human populations, rather than only reflecting and benefiting those in industrialized nations, it is necessary to involve indigenous populations in research in a legal, ethical, and non-exploitative manner (Abdill et al., 2022; Green et al., 2020). Here, we isolated live bacterial strains from anonymized fecal samples collected from Hadza hunter-gatherers in 2013/2014 (Fragiadakis et al., 2019; Merrill et al., 2022; Smits et al., 2017). Samples were collected with permission from the Tanzanian government, National Institute of Medical Research (MR/53i 100/83, NIMR/HQ/R.8a/Vol.IX/1542), the Tanzania Commission for Science and Technology, and with aid from Tanzanian scientists. A material transfer agreement with the National Institute for Medical Research in Tanzania specifies that collected samples are solely to be used for academic purposes. For more information on the consent practices followed, and our ongoing work to communicate the results of these projects to the Hadza, please see (Merrill et al., 2022; Olm et al., 2022).

## Introduction

The industrialized lifestyle is defined by the consumption of highly-processed foods, high rates of antibiotic administration, cesarean section births, sanitation of the living environment, and reduced contact with animals and soil–all of which can impact the human gut microbiota (Sonnenburg and Sonnenburg, 2019). Certain taxa are influenced by industrialization, i.e., are prevalent and abundant in non-industrialized populations and diminished or absent in industrialized populations, or vice versa. (De Filippo et al., 2010; Jha et al., 2018; Merrill et al., 2022; Olm et al., 2022; Smits et al., 2017; Vangay et al., 2018; Yatsunenko et al., 2012). The microbiome of 1000-2000 year-old North American paleofeces is more similar to the modern non-industrialized than industrialized gut (Wibowo et al., 2021). The industrialized microbiota appears to be a product of both microbial extinction, as once-dominant taxa disappear, and expansion of less dominant or new taxa (Sonnenburg and Sonnenburg, 2014).

The industrialized diet differs drastically from non-industrialized diets, including in reduced amount of microbiota-accessible carbohydrates (MACs), a major metabolic input for microbes in the distal gastrointestinal tract (Cordain et al., 2005; Flint et al., 2012; Sonnenburg and Sonnenburg, 2014). Some gut-resident microbes use host mucin, which is heavily glycosylated, as a carbon source, depending on the availability of dietary MACs (Bell and Juge, 2021; Desai et al., 2016; Pudlo et al., 2022; Salyers et al., 1977; Sonnenburg et al., 2005). Shifts in dietary MACs alter microbial relative abundances and may increase inflammation and susceptibility to intestinal pathogens (Desai et al., 2016; Earle et al., 2015; Martens et al., 2018). Taxa are lost due to a lack of dietary MACs over generations in a mouse model, (Sonnenburg et al., 2016) and in humans as they immigrate to the U.S. (Vangay et al., 2018).

As human populations adopt an industrialized lifestyle, the prevalence of *Prevotella* decreases and that of *Bacteroides* increases (De Filippo et al., 2010; Jha et al., 2018; Kaplan et al., 2019). While *Bacteroides* are well-studied, *Prevotella* species remain understudied with few tools available for mechanistic investigation (Abdill et al., 2022; Accetto and Avguštin, 2015; Li et al., 2021; Xu et al., 2003). Both genera harbor well-documented carbohydrate utilization capabilities, encoded in carbohydrate active enzymes (CAZymes), often organized into polysaccharide utilization loci (PULs) (Bjursell et al., 2006; Dodd et al., 2010; Fehlner-Peach et al., 2019). Characterization of intestinal *Prevotella* species have been limited by challenges with colonization, particularly mono-colonization of germ-free mice. Here we overcome these barriers to establish a causal link between diet and *P. copri* abundance in a gnotobiotic mouse model.

The decreased prevalence of *Prevotella* in industrial populations is likely linked to a decline in relative abundance within individual microbiomes (Sprockett et al., 2020). Decreased abundance of bacterial taxa in individuals reduces the likelihood of transmission from mother to infant (Olm et al., 2022; Sonnenburg and Sonnenburg, 2019). When compounded over generations, decreased abundance can result in a population-level decline in prevalence and eventually taxa loss or extinction (Vangay et al., 2018; Sonnenburg et al., 2016). The factors driving the decline in *Prevotella* and the increase in *Bacteroides* during industrialization remain to be defined. The abundance and prevalence of specific strains of *P. copri*, the dominant *Prevotella* species in the human gut, vary among populations based on host lifestyle, particularly diet (De Filippis et al., 2019; Tett et al., 2019). Here we use gnotobiotic mice to investigate the role of diet in sustaining Prevotella and Bacteroides colonization; we demonstrate that dietary MACs play a key role in controlling the abundances of *Bacteroides* and *Prevotella*.

## Results

### *Bacteroides* and *Prevotella* genomes from the Hadza microbiota vary in prevalence across populations

To compare *Prevotella* and *Bacteroides* from non-industrialized lifestyle populations, we isolated and sequenced 6 *Bacteroides* strains (from 4 species) and 7 *Prevotella copri* strains from stool samples collected from 13 Hadza individuals. Importantly, *P. copri* is a species known to encompass extensive genomic diversity and its division into multiple species has been discussed (Tett et al., 2019). Single isolate genomes were assembled using both MiSeq generated short reads (146bp) and nanopore generated long reads (10-100kb) (<u>Table 1</u>).

To determine relatedness of the Hadza genomes to previously sequenced genomes, we calculated the average nucleotide identity (ANI) distance to the closest genome present in the National Center for Biotechnology Information (NCBI) GenBank. The Hadza *Prevotella* genomes are statistically distinct from *P. copri* genomes classified as complete in NCBI (p=0.0002; Wilcoxon rank-sums test). The Hadza *Bacteroides* genomes, however, are not statistically distinct from existing complete *Bacteroides* genomes in NCBI (p=0.76; Wilcoxon rank-sums test). The difference in distinctness of the Hadza *Prevotella* and *Bacteroides* genomes is unlikely due to an underrepresentation of *Prevotella* genomes as each genus has a similar number of published genomes (3482 and 4199, *Prevotella* and *Bacteroides*, respectively). It is also worth noting that the vast majority of human gut microbiota genome sequences were collected from North American and European samples (Abdill et al., 2022).

A phylogenetic comparison of the sequenced Hadza strains to representative genomes of the same species reveals that the Hadza *Prevotella* and *Bacteroides* strains cluster with, but are distinct from, type strains and other respective species (**Fig. 1A, Fig. S1**). All 7 Hadza *P. copri* strains belong to Clade A of the 4 proposed *P. copri* subgroups possessing >10% inter-clade genetic divergence (Tett et al., 2019).

**Figure 1.**
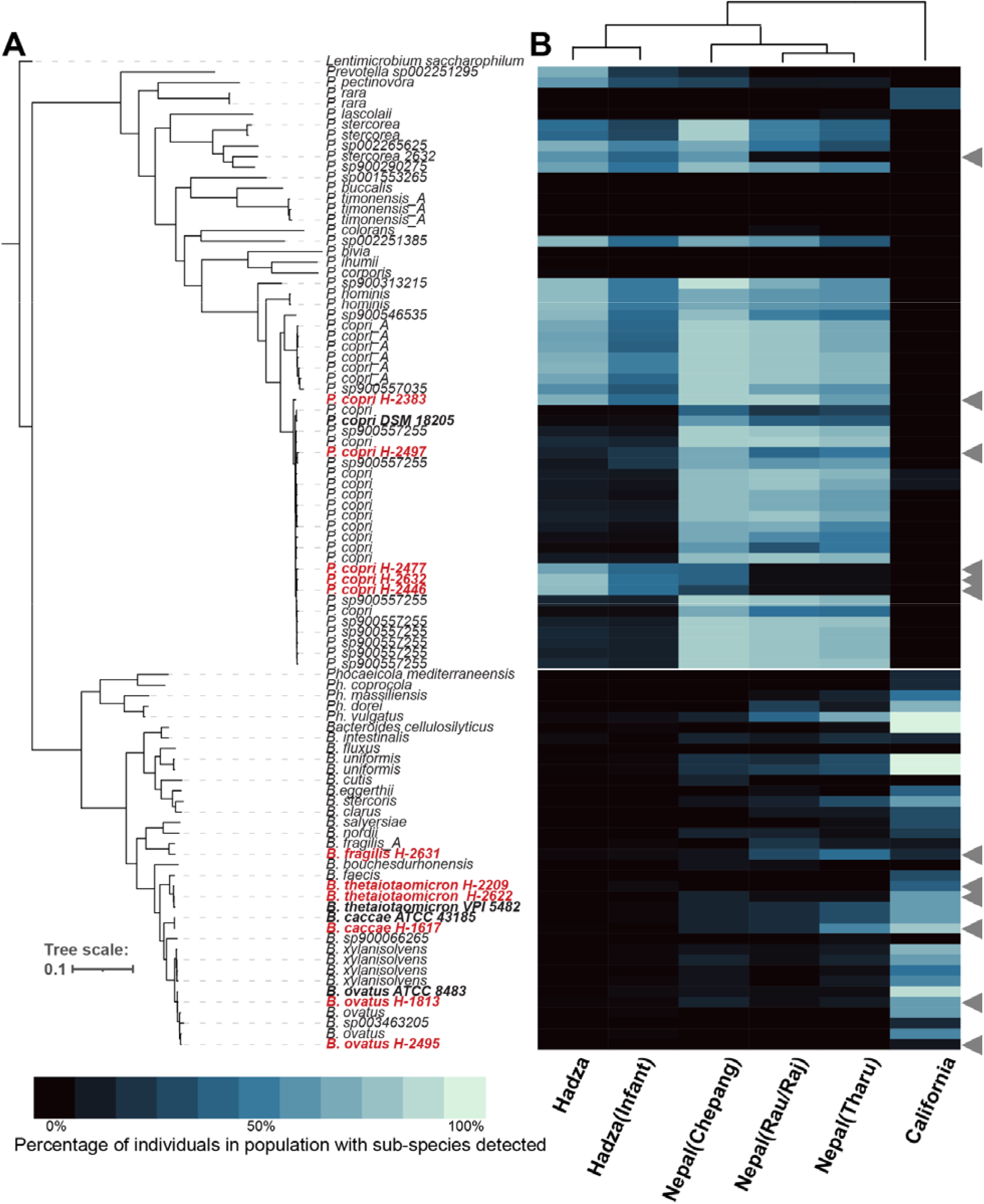
Hadza *Bacteroides* and *Prevotella* strains are related to previously sequenced isolates and vary in prevalence across populations. **(A)** Phylogenetic tree of *Prevotella* and *Bacteroides* genomes. Isolates from this study (red) and genomes from GenBank (black), strains used later in this study (bold). **(B)** *Prevotella* and *Bacteroides* subspecies prevalence across foragers (Hadza and Chepang), agriculturalist (Rau, Raj, Tharu), or industrialized (California). Prevalence defined as % of gut metagenomes from a population (column) in which particular strain (row) is detected.

To understand the prevalence of these genomes across human populations, we compared *Prevotella* and *Bacteroides* prevalence among Hadza adults and infants, four populations from Nepal living on a lifestyle gradient including foraging (Chepang), recent agriculturalist (Raute, Raj), longer term agriculturalist (Tharu), and industrial lifestyle populations (California) (**Fig. 1B)**. We chose these groups due to their varied lifestyles and the exceptional metagenomic sequencing depth achieved, averaging 23 Gbp per sample (Jha et al., 2018; Merrill et al., 2022). Of the populations analyzed, the prevalence of Hadza *Prevotella* and *Bacteroides* isolate genomes are most similar to another foraging group, the Chepang, and most distinct from industrial lifestyle individuals (California). Prevotella genomes are rare or absent from the industrialized populations, while more prevalent and abundant in the Hadza and agriculturist samples. Conversely, nearly all Bacteroides genomes, including those isolated from the Hadza, are more prevalent in industrialized populations. The clear lifestyle shift associated with *Bacteroides* and *Prevotella* prevalence leads to the question of what aspects of the industrial lifestyle have driven these changes.

### Dietary MACs are necessary for *P. copri* persistence

While many factors differentiate the industrial and non-industrial lifestyles, diet serves as the top candidate for driving microbiome alterations (Sonnenburg and Sonnenburg, 2014). The Hadza diet is rich in dietary MACs from foraged tubers, berries, and baobab (Marlowe and Berbesque, 2009). In contrast, the industrialized diet is typified by high caloric intake and foods rich in fat and low in MACs (Monteiro et al., 2013). We wondered whether diet alone could impact the ability of Hadza *Bacteroides* and *Prevotella* to colonize mice. Germ-free (GF) mice were colonized with either Hadza *B. thetaiotaomicron (Bt) H-2622*, or Hadza *P. copri (Pc) H-2477*. Mice were maintained on a high MAC diet for 7 days and then switched to either a diet devoid of MACs (no MAC), a high-fat/low-MAC diet (Western), or maintained on the high MAC diet for 7 days (**Fig. 2A**). *Bt H-2622* colonization density (10^9^ CFU/ml in feces) on the high MAC diet was maintained in all three diet conditions (**Fig. 2B**). *Pc H-2477* colonized to a lower degree on the high MAC diet (10^7^ CFU/ml) and declined drastically following the change to the Western or no MAC diet, with no fecal CFUs detectable 7 days post diet switch (**Fig. 2C**). The lack of detectable *Pc H-2477* in the absence of dietary MACs was particularly striking given the absence of competition from other microbes in this mono-associated state. To our knowledge this is the first example of a strain’s apparent eradication in a mono-associated state due to a diet change. Two other *P. copri* strains (Hadza *Pc H-2497* and a non-Hadza strain isolated from an individual of African origin *Pc N-01*) are also lost *in vivo* in the absence of dietary MACs (**Fig. S2 A, B**), indicating that survival of *P. copri in vivo* depends on the presence of dietary MACs.

**Figure 2.**
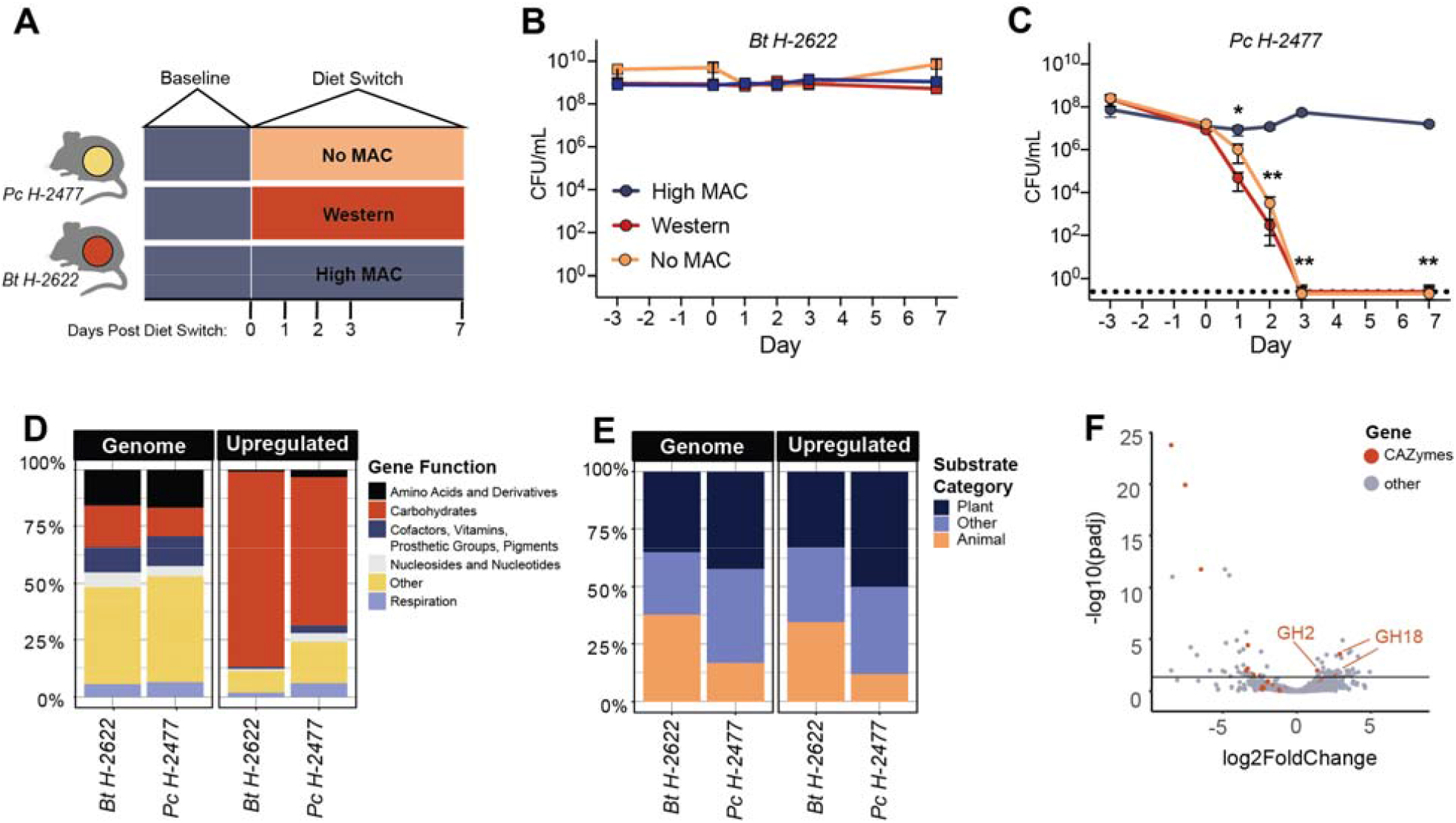
*Bt* and *Pc* colonization differ in diet-dependent manner. **(A)** Schematic of gnotobiotic experiments. **(B, C)** Fecal density of *Bt H-2622* (B), and *Pc H-2477* **(C)**, in monocolonized mice (n=4/group for High MAC and Western conditions, n=5/group for No MAC conditions) fed different diets (multiple Mann-Whitney tests, *: p ≤ 0.05, **: p ≤ 0.01). Representative experiments, repeated twice. Dotted line denotes 0 CFU. **(D)** Proportion of genes upregulated *in vivo* by functional categories (RAST). **(E)** Proportion of predicted substrate categories of upregulated CAZymes. **(F)** Differential expression of *Bt H-2622* genes in No MAC versus High MAC diets.

To better understand the strategies used by Hadza *Pc* and *Bt* to persist *in vivo*, we analyzed transcriptional profiling data from cecal contents of mice monocolonized with either *Pc H-2477* or *Bt H-2622* fed a high MAC diet relative to *in vitro* growth in peptone yeast glucose broth (PYG). *Bt H-2622* and *Pc H-2477* upregulate many genes *in vivo* under high MAC diet conditions (**Fig. S2C, D**). Despite the fact that 18% and 13% of genes in *Bt H-2622* and *Pc H-2477*, respectively, encode for predicted carbohydrate utilization proteins, 86% (in *Bt H-2622*) and 65% (in *Pc H-2477*) of those upregulated *in vivo* encode for carbohydrate utilization (p < 3.7e-12 for *Bt*, p < 4.5e-13 for *Pc*, Fisher’s Exact test), indicating that carbohydrate utilization is the major metabolic function of these organisms *in vivo* (**Fig. 2D**).

A comparison of glycosidic linkage-breaking CAZymes, glycoside hydrolases (GH) and polysaccharide lyases (PL), reveals that *Bt H-2622* upregulates a higher proportion of GHs and PLs devoted to animal-derived carbohydrate utilization relative to *Pc H-2477* (**Fig. 2E**). Specifically, *in vivo* under high MAC diet conditions *Bt H-2622* upregulates 8 of 22 encoded mucus targeted GHs (3/10 GH18; 5/12 GH20) whereas *Pc H-2477* encodes no GH18s and only one GH20, which is not upregulated in the high MAC diet condition. In addition to targeting mucus carbohydrates, *Bt H-2622* also upregulates 40 of its 97 plant-targeting GHs and PLs whereas *Pc H-2477* upregulates all 38 of its plant-targeting GHs and PLs in the high MAC diet (**Fig. 2E**). On the no MAC diet, *Bt H-2622* upregulates 2 additional GH20s (along with the other mucin CAZymes upregulated on the high MAC diet) as well as 27 plant-targeting GH and PLs relative to the *in vitro* condition (**Fig. S2E**). When comparing the high MAC and no MAC *in vivo* conditions, *Bt H-2622* upregulates only 3 GHs, 2 of which degrade mucin (GH18) (**Fig. 2F**) (Sonnenburg et al., 2005). These data indicate that in the absence of diet derived MACs, Hadza *Bt H-2622* relies on mucus carbohydrates and that limited mucin degrading capabilities render *Pc H-2477* incapable of sustaining colonization in the absence of dietary MACs.

### Carbohydrate degradation capacity differs between Hadza *Bacteroides* and *Prevotella* mirrors industrialized strains

Hadza *Pc* and *Bacteroides* isolates have a similar number and predicted function of GHs and PLs to reference strains of the corresponding species (<u>Table 2</u>). Unsupervised clustering of GHs and PLs reveals that the Hadza strains cluster with their type strain counterparts (**Fig. 3A**). When comparing the total number of GHs and PLs encoded within the Hadza strains to non Hadza strains, we found similar total numbers of these genes and distribution of substrate specificity between strains of the same species (**Fig. 3B**).

**Figure 3.**
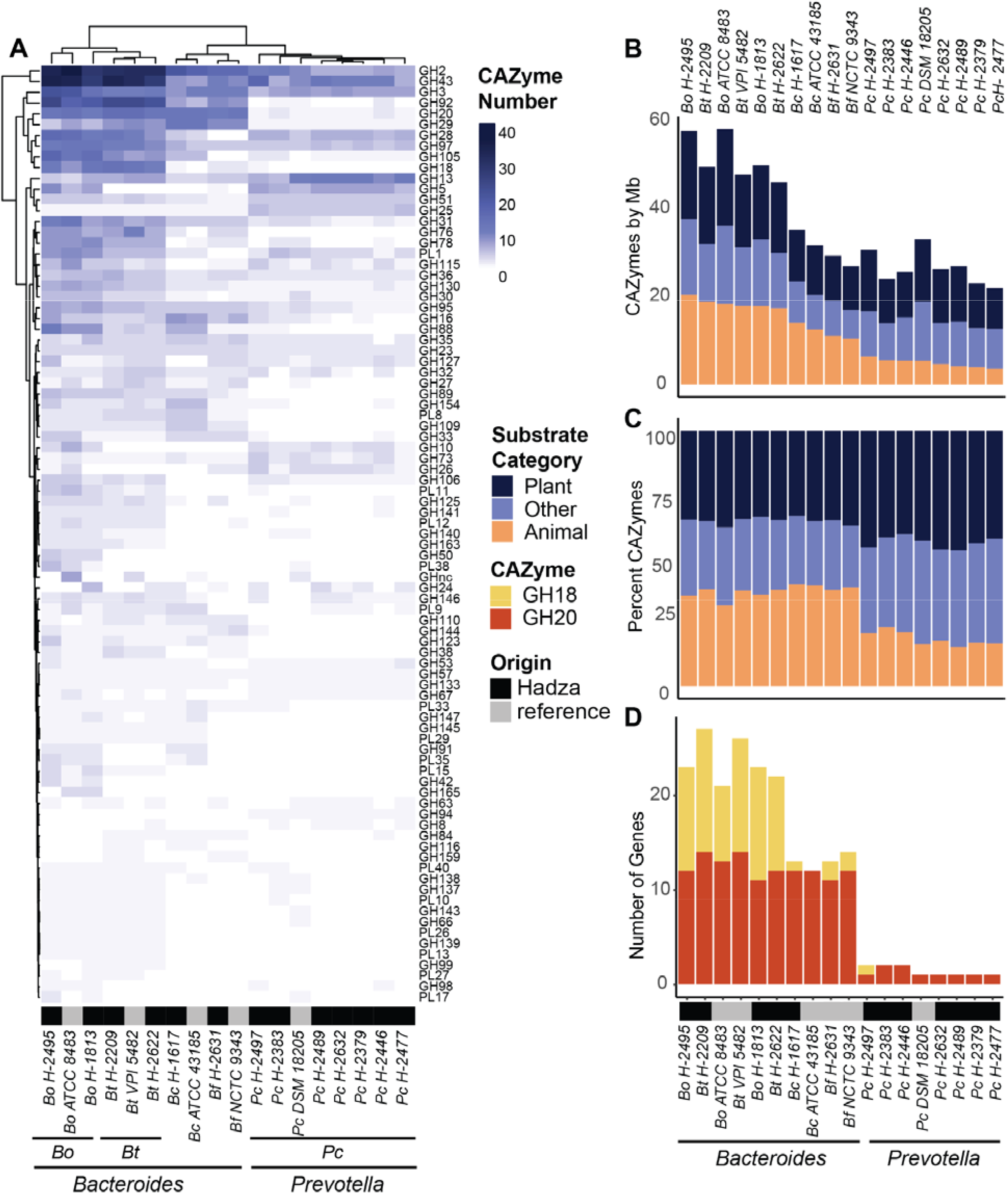
Hadza *Bacteroides* and *Prevotella* differ in distribution of glycoside hydrolases (GH) and polysaccharide lyases (PL). **(A)** Number of GHs and PLs per genome indicated by CAZy family (rows); CAZy families are shown if represented in at least 4 genomes analyzed. **(B)** Number of GHs and PLs normalized to genome size (Mb), colored by predicted substrate. **(C)** Proportion of GHs and PLs in each genome colored by predicted substrate. **(D)** Number of mucin-degrading GH18 and GH20 genes per genome.

While Hadza *Bacteroides* and *Prevotella* strains mirror the carbohydrate degrading capacity of their non Hadza counterparts, large differences exist between the *Bacteroides* and *Prevotella* strains. The *Bacteroides* encode more GHs and PLs than *Prevotella* strains even when corrected for genome size (251/21 average GH/PL in *Bacteroides*; 101/5 in *Prevotella*; Welch Two Sample t-test p = 0.00561) **(Table 2, Fig. 3B)**. The proportion of *Bacteroides* GHs and PLs that are predicted to target plant carbohydrates or animal carbohydrates are equivalent (average 34% and 37%, respectively) whereas the *Prevotella*-encoded carbohydrate degradation is biased toward plant over animal carbohydrates (average 44% and 19%, respectively) **(Fig. 3C)**. The *Bacteroides* also encode a greater breadth of GH and PL families (averaging 68 CAZyme families per genome) while *Pc* isolates average 40 CAZy families per genome (**Fig. S3A**), consistent with previously reported distributions for industrial lifestyle derived *Bacteroides* and *Prevotella* strains (Fehlner-Peach et al., 2019). The two genera also differ in their predicted mucin-degradation capacity. CAZyme families GH18 and GH20 target carbohydrates found within the intestinal mucus lining (Luis et al., 2021). All Hadza *Bacteroides* isolates harbor 11-14 GH20 and 1-13 GH18 CAZymes, however the Hadza *Prevotella* isolates contain only 1 or 2 GH20s and only one isolate, *Pc H-2497*, contains a single GH18 **(Fig. 3D, S3B)**.

The CAZyme content of Hadza *Bacteroides* and *Prevotella* isolates are similar to their non-Hadza counterparts. Hadza *Bacteroides* isolates contain both more GHs and PLs overall as well as broader substrate degrading capabilities that include both plant and animal derived carbohydrates relative to the Hadza *Prevotella* isolates. This difference between the Hadza *Bacteroides* and *Prevotella* strains is similar to that seen in non-Hazda strains suggesting that the *Prevotella* niche is more reliant upon plant carbohydrates compared to *Bacteroides* (Gálvez et al., 2020).

### Dietary MACs are sufficient to maintain *Pc* colonization in the presence of *Bt*

To test whether Hadza *Bacteroides* and *Prevotella* isolates differ in their ability to use plant and mucus derived carbohydrates, we cultured Hadza and type strain *Bacteroides* and *Pc* isolates in media containing the plant carbohydrate inulin, porcine gastric mucin glycans, porcine intestinal heparin, or fructose as the sole carbon source. There is a range of ability to utilize inulin across the strains, consistent with previous work (**Fig. 4A**) (Sonnenburg et al., 2010). Growth in the presence of mucin, however, is divided by genera; most *Bacteroides* isolates grow well on mucin but the *P. copri* isolates do not **(Fig. 4A)**. These data are consistent with the lack of mucin degrading capacity within the *Pc* genomes and the loss of *Pc* colonization *in vivo* when the host is the sole carbohydrate source.

**Figure 4.**
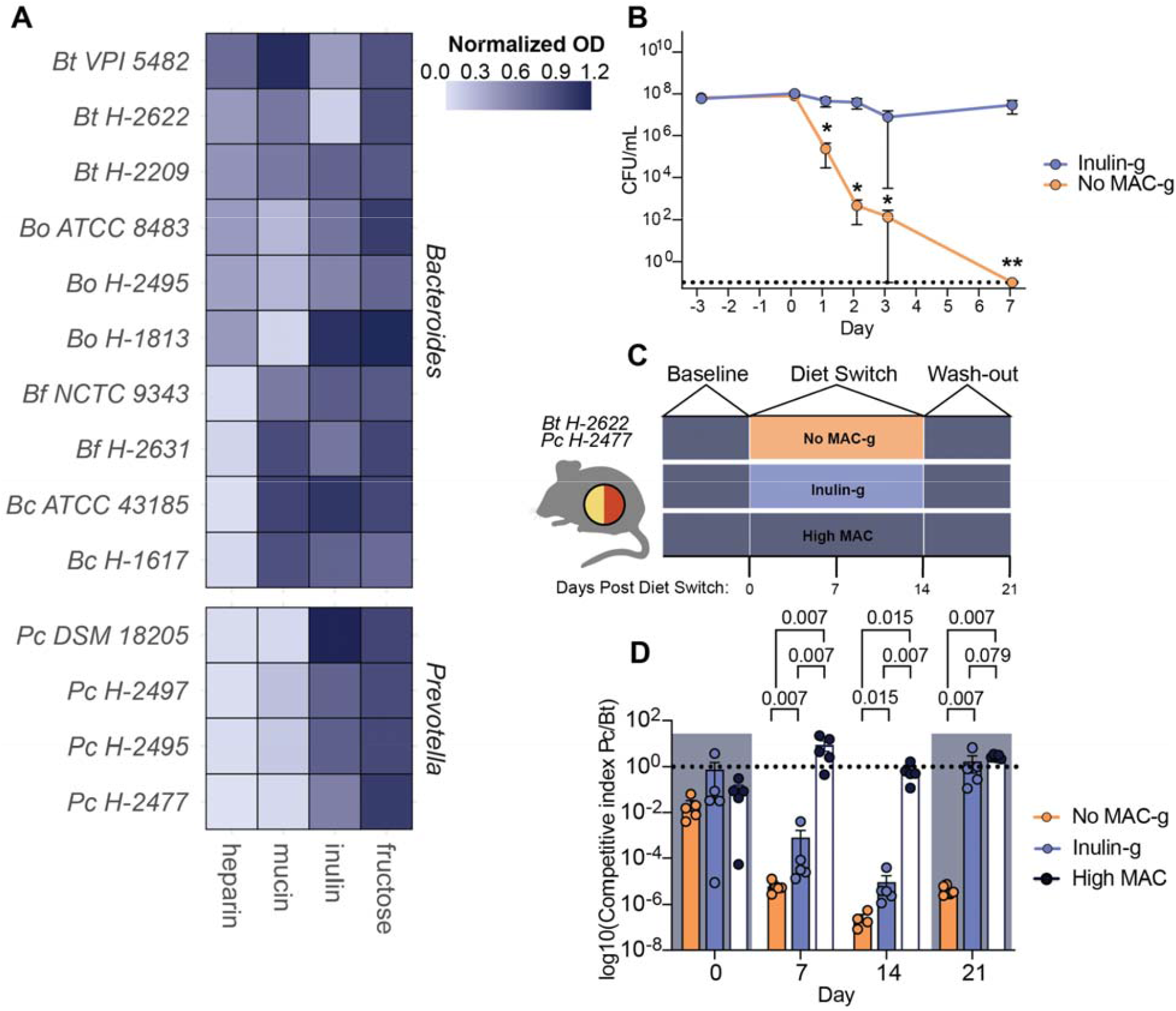
Reintroduction of MACs is sufficient to maintain *P. copri* colonization. **(A)** Normalized maximum OD600 of *Bacteroides* and *Prevotella* isolates grown in YCFA with a single added carbohydrate for 24 hours. **(B)** Fecal CFUs of *Pc H-2477* in monocolonized mice fed a No MAC-g or Inulin-g diet (mean + s.e.m., n=5 mice per group, Multiple Mann-Whitney tests, *: p ≤ 0.05, **: p ≤ 0.01). Dotted line denotes 0 CFU. Representative experiment, repeated three times. **(C)** Schematic of bicolonization with *Pc H-2477* and *Bt H-2622*. **(D)** Ratio of *Pc H-2477* to *Bt H-2622* (mean + s.e.m., n= 5 mice per group, K-S test). Dotted line denotes equal abundance. Shading indicates administration of High MAC diet. Representative experiment, repeated twice.

To determine whether the lack of diet-derived MACs is responsible for the loss of *Pc H-2477* colonization *in vivo*, we fed mice mono-colonized with *Pc H-2477* a high MAC diet and then switched to either a custom diet containing 34% inulin by weight as the sole fermentable carbohydrate to match MAC content of the high MAC diet (custom diets use gelatin as a binding agent and are noted by a “-g”; Inulin-g) or a no MAC diet (no MAC-g) (**Fig. 4B)** (Dubos and Pierce, 1948). The no MAC-g diet did not sustain *Pc H-2477* colonization, with the strain becoming undetectable within one week (**Fig. 4B**). However, *Pc H-2477* maintained colonization in the presence of the Inulin-g diet to levels similar to those observed in the high MAC diet (**Fig. 2C, 4B**), consistent with the requirement of dietary MACs for *Pc H-2477* colonization *in vivo*.

We were curious how dietary MACs impact the relative abundance of *Pc* and *Bt* in mice when colonized together. GF mice were co-colonized with *Pc H-2477* and *Bt H-2622* and fed a high MAC diet for 7 days and then either maintained on the high MAC diet, switched to the no MAC-g diet, or the Inulin-g diet for 2 weeks, followed by a one week period in which all mice consumed the high MAC diet (**Fig. 4C**). Prior to the diet switch (Day 0), mice harbored both *Pc H-2477* and *Bt H-2622*. However, 7 days after the switch to either the no MAC-g, *Pc H-2477* decreased dramatically in abundance relative to *Bt H-2622*; a decrease of *Pc* also occurred in the Inulin-g diet, however the drop was not as severe as the no MAC diet indicating that inulin provided support to this strain (**Fig 4D**). When mice were returned to the high MAC diet, those fed the Inulin-g diet regained relative abundance of *Pc H-2477* equivalent to that of baseline and to mice fed the high MAC diet throughout the experiment (**Fig. 4E**). In mice switched to the high MAC diet from the no MAC-g diet, *Pc H-2477* colonization was detectable, but remained low after 7 days on the high MAC diet. These data are consistent with the requirement of dietary MACs for *Pc* colonization in the presence of *Bt* and may show that the variety of carbohydrates in the high MAC diet (derived from wheat, corn, oats, and alfalfa) better supports *Pc* colonization than a single MAC source like inulin under competition from Bacteroides. Furthermore, prolonged absence of dietary MACs restricts the ability of *Pc* to regain abundance when MACs are reintroduced.

## Discussion

The tradeoff between a microbiome dominated by *Bacteroides* or *Prevotella* based on host lifestyle has been well described, but its basis is not well understood (Gorvitovskaia et al., 2016; Yatsunenko et al., 2012). Here we demonstrate that Hadza isolates of *Bacteroides* and *Prevotella* do not differ dramatically from their non-Hazda counterparts in terms of genome-wide average nucleotide identity and carbohydrate utilization, suggesting that differences in their relative abundance and prevalence across lifestyle is not due to an inherent property of the population-specific strains themselves, but to differences in their environments. Furthermore, we demonstrate that dietary MACs are crucial for *Prevotella* to maintain colonization: even as the sole microbe, *Prevotella* is eradicated when dietary MACs are removed. *Bacteroides* species, however, can maintain colonization in the absence of dietary MACs due to their ability to use both plant- and host-derived carbohydrates, enabling continued colonization in low MAC industrialized diets. Our data demonstrates that in the presence of dietary MACs in gnotobiotic models, Hadza *Bacteroides* and *Prevotella* can co-exist, as is seen in the Hadza microbiome. However removal of dietary MACs results in a precipitous decline in *Prevotella*, which is slow to recover when MACs are reintroduced. The presence of a single MAC in the diet, inulin, was sufficient to maintain an intermediate level of colonization that then rebounded when a more complete palate of MACs was available. These data are reminiscent of the seasonal pattern of *Prevotella* abundance in the Hadza, which cycles in abundance with the seasonality of their diet.

All together these data are consistent with the model that prior to industrialization, human microbiomes harbored both *Bacteroides* and *Prevotella* species. As diets shift from high MAC foraged foods to low MAC industrially produced foods, abundance and prevalence of *Prevotella* diminished to the point of extinction in some individuals (Merrill et al., 2022). How the loss of *Prevotella* and increased abundance of *Bacteroides* within the industrialized microbiome impacts human physiology remains an important question.

## Acknowledgements

We thank Bryan Merrill, Madeline Topf, and Michelle St. Onge for technical support; Gabriela Gall Rosa for help with analysis; Samuel Lancaster, Brittany Sison, and Audrey Zhang for assistance processing RNAseq data; Michael Fischbach and Niokhor Dione for providing *P. copri N-01*; Brian Yu and Rose Yan for sequencing support; David Schneider for advice and tissue culture hood use; Bernard Henrissat for advice on CAZyme annotation.

R.H.G. was supported by an NIH/NHGRI T32 training grant (5T32HG000044-23) and a Blavatnik Family Fellowship. This work was supported by grants from the NIH to J.L.S. (R01-DK085025, DP1-AT009892). J.L.S. is a Chan-Zuckerberg Biohub investigator.

## Author Contribution

R.H.G., M.R.O., N.T., F.E., and S.K.H. performed the experiments and designed and performed data analyses. R.H.G., J.L.S., and E.D.S. conceived of the study and wrote the manuscript.

## Disclosure Statement

J.L.S. is a co-founder, shareholder, and on the scientific advisory board of Novome Biotechnologies and Interface Biosciences.

## Star Methods

### KEY RESOURCES TABLE

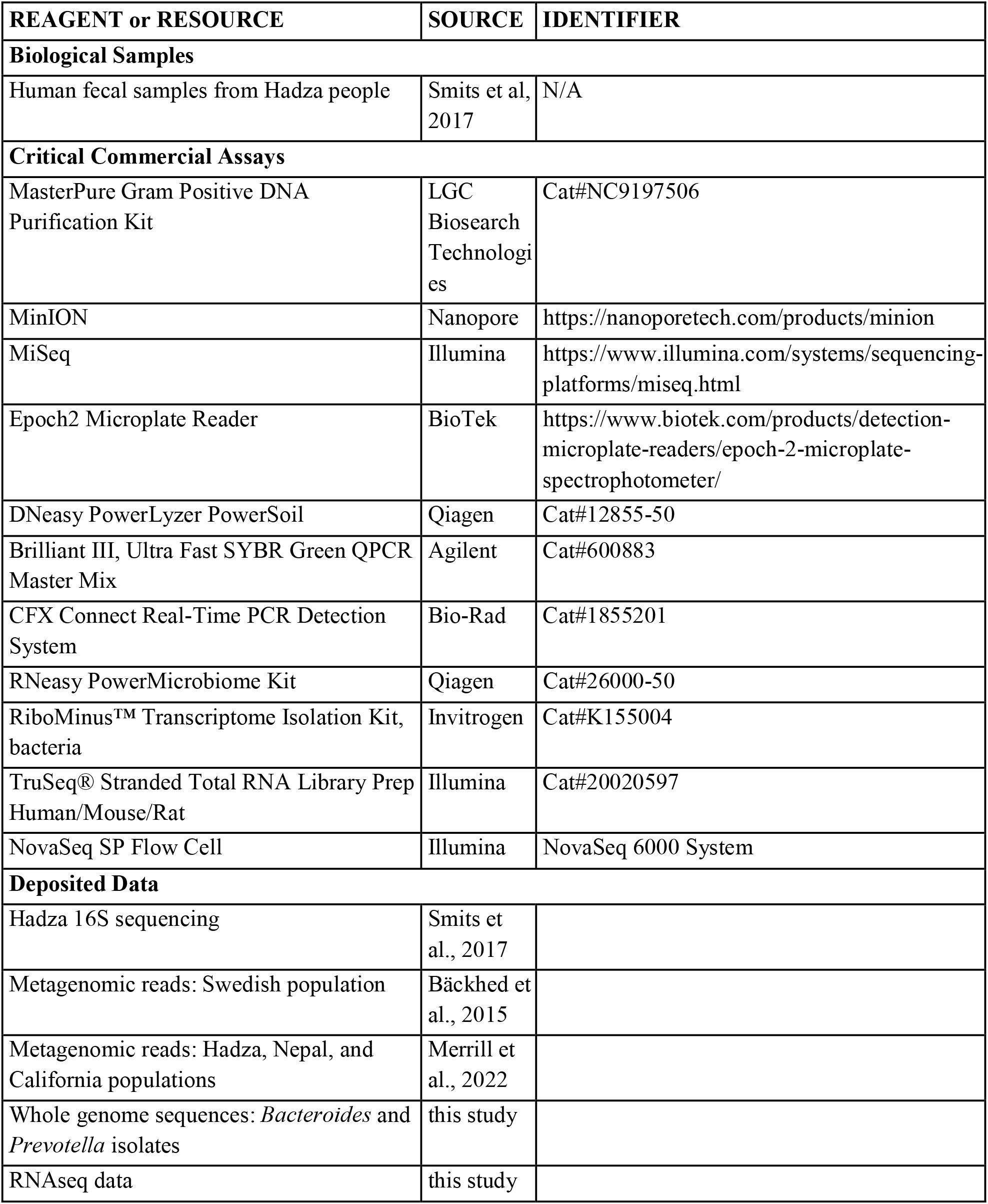
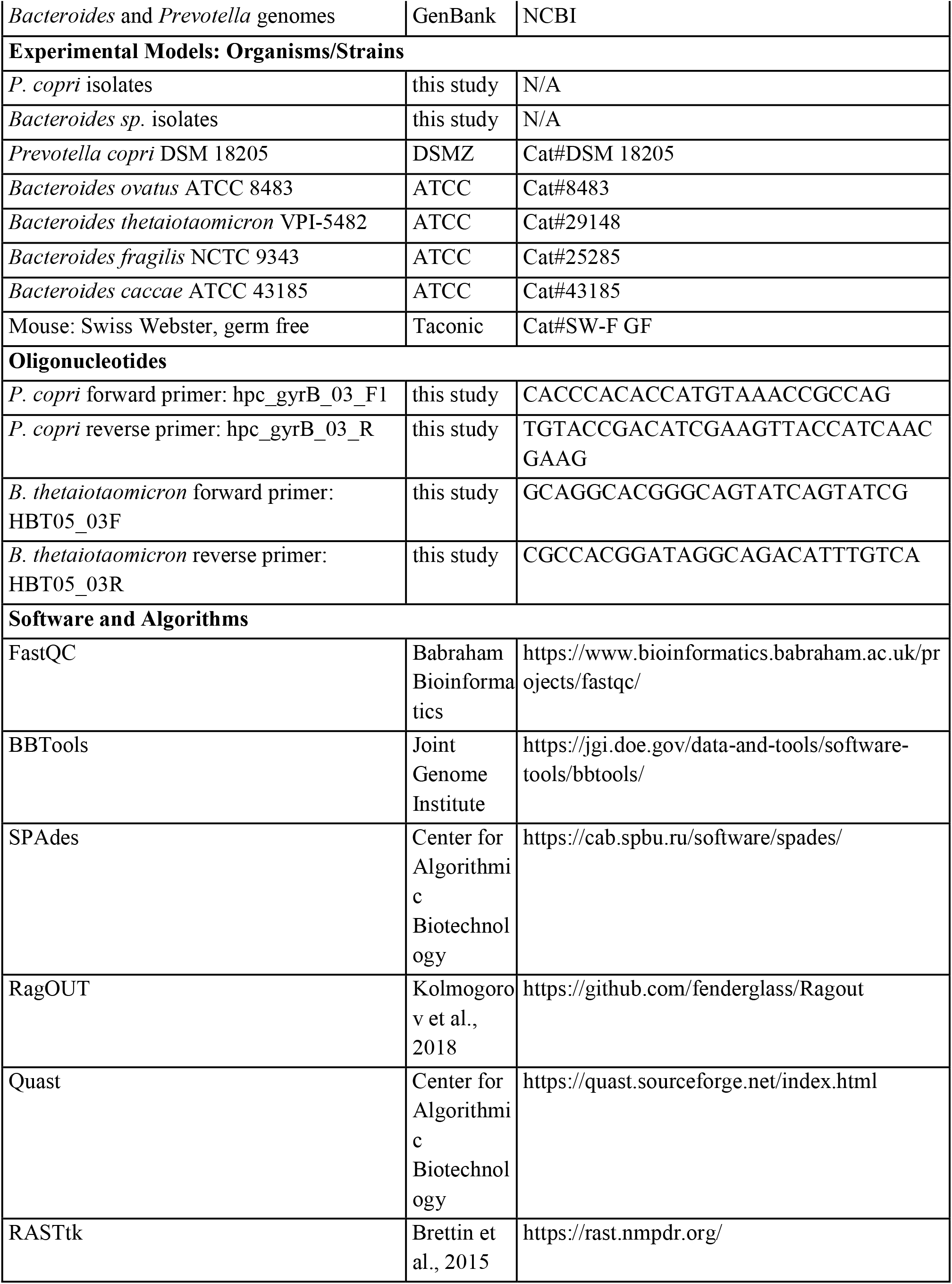
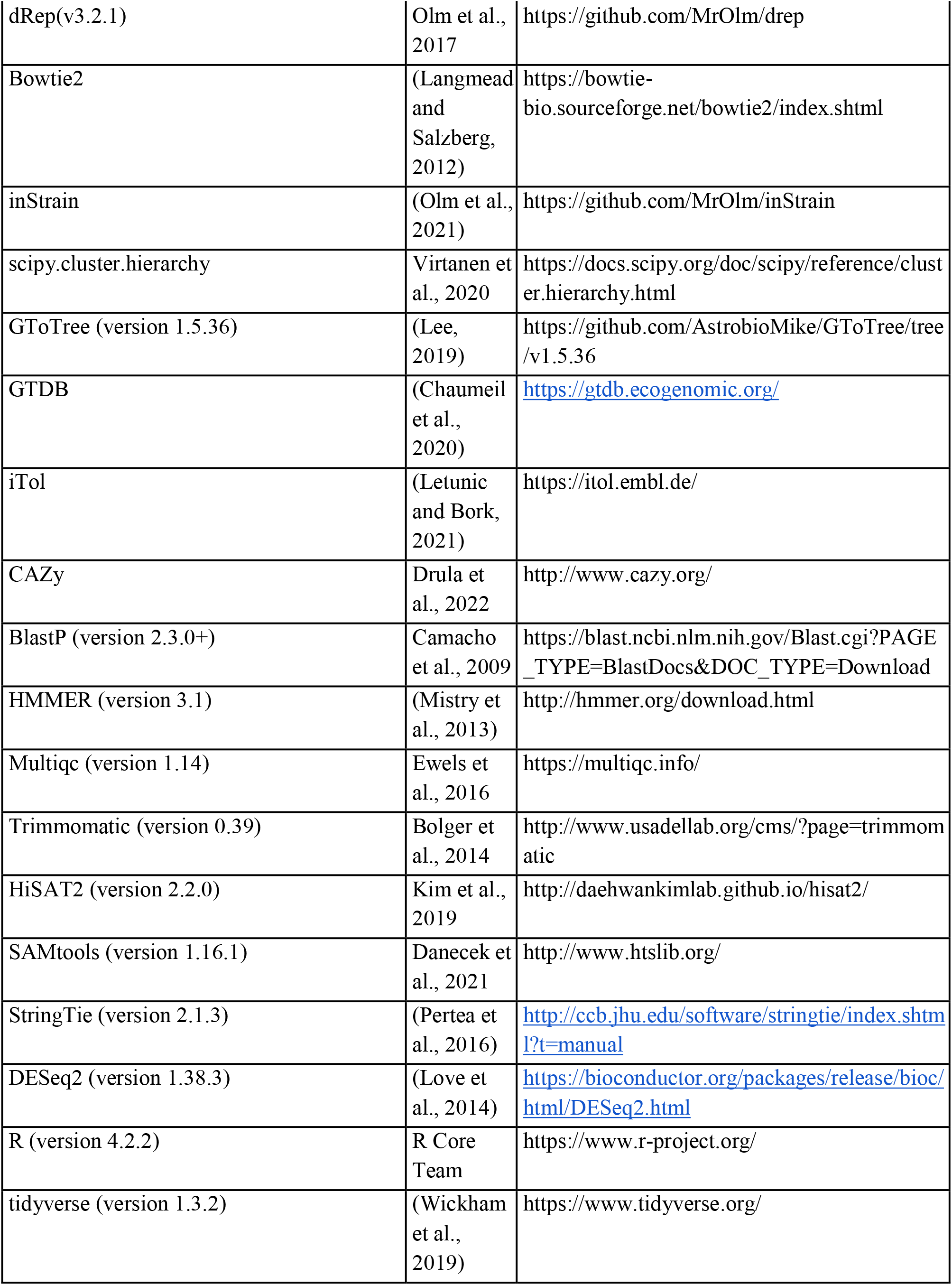
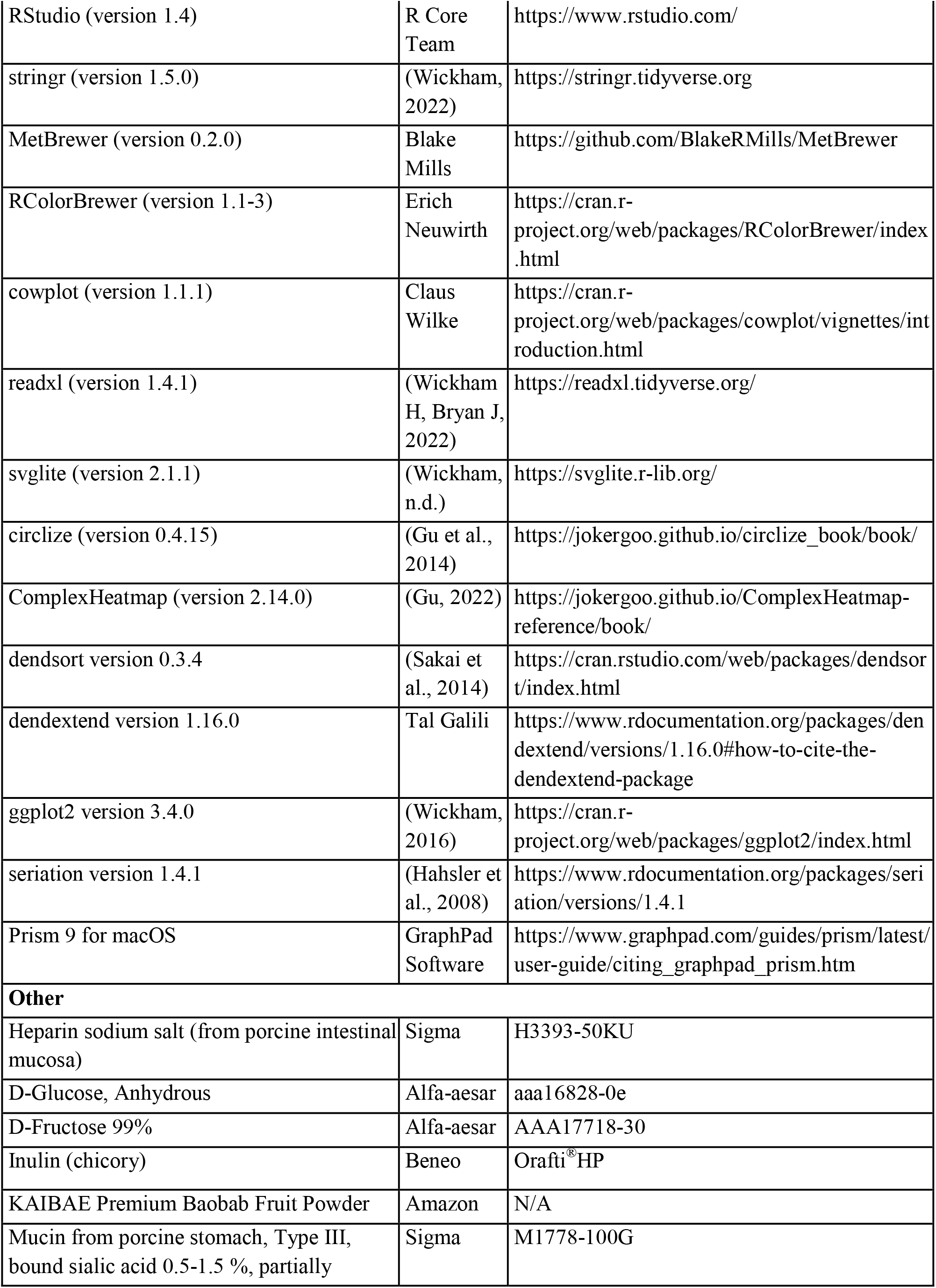
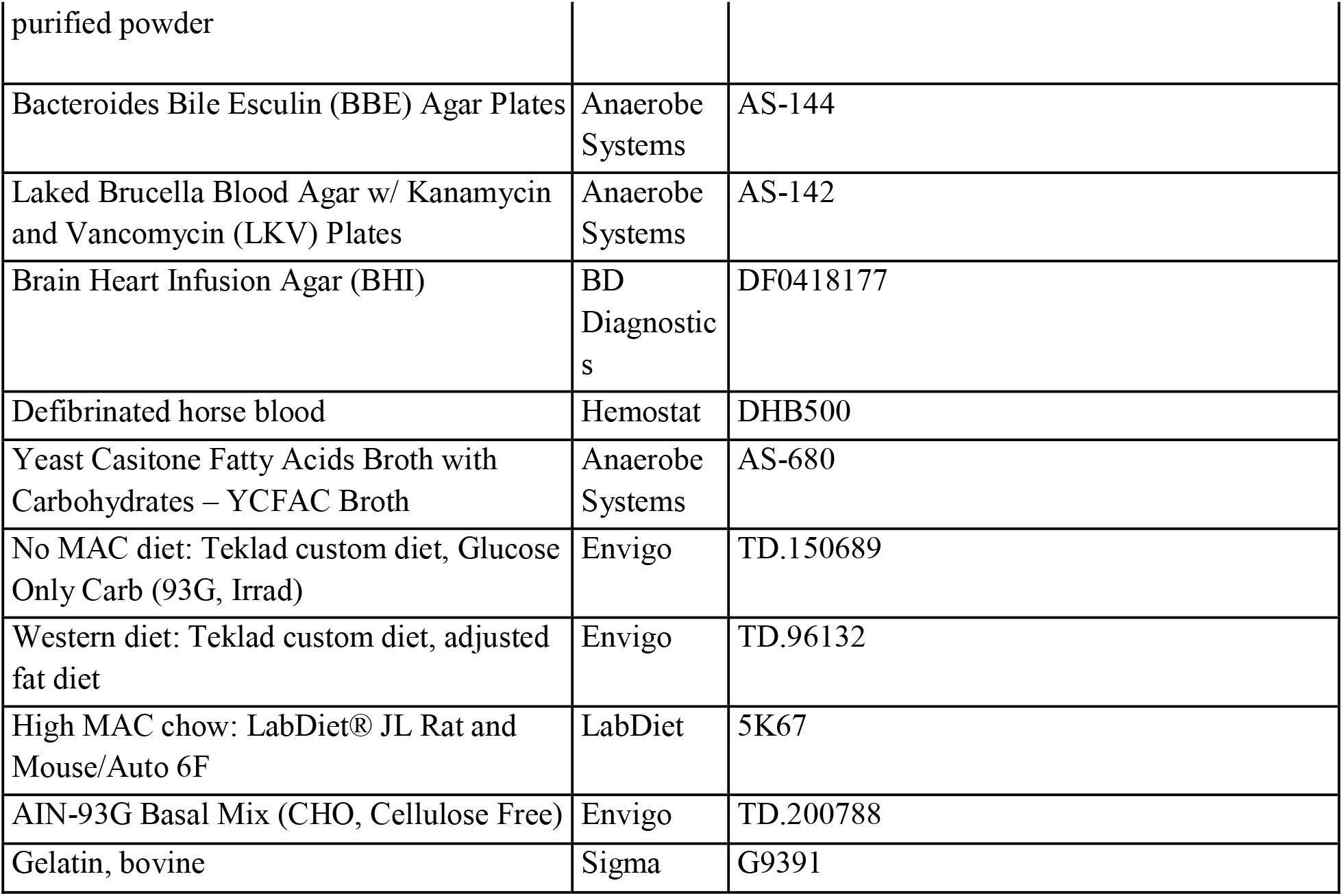

#### LEAD CONTACT AND MATERIALS AVAILABILITY

All information and requests for further resources should be directed to and will be fulfilled by the Lead Contact, Erica Sonnenburg

### Data and code availability

Datasets and code for analysis are available at https://github.com/SonnenburgLab/. Raw data files for WGS are in the process of being uploaded to public databases and will be freely available upon publication of this manuscript.

## Experimental Model Details

### Bacterial Culture

Bacteria not isolated in this study were purchased from DSMZ (*P. copri DSM 18205*), or ATCC (all other reference strains). Glycerol stocks were struck out on Brain Heart Infusion plates with 10% defibrinated horse blood (BHIBA) and incubated anaerobically for 24-48 h at 37ºC. All growth and culturing of *Bacteroides* and *Prevotella* strains were performed anaerobically in a Coy anaerobic chamber containing 87% N2, 10% CO2, and 3% H2.

### Mouse Husbandry

All mouse experiments were performed in accordance with the Stanford Institutional Animal Care and Use Committee. Mice were maintained on a 12-h light/dark cycle at 20.5 □ °C at ambient humidity, fed ad libitum, and maintained in flexible film gnotobiotic isolators for the duration of all experiments (Class Biologically Clean). Swiss-Webster mice were used for gnotobiotic experiments and the sterility of germ-free mice was verified by 16S PCR amplification and anaerobic culture of feces. Sample sizes were chosen on the basis of litter numbers and controlled for sex and age within experiments. Researchers were unblinded during sample collection (Pruss and Sonnenburg, 2021).

## Method Details

### Strain Isolation from Fecal Samples

Samples for strain isolation were chosen from the samples reported previously based on the 16S abundance of either *Bacteroides* or *Prevotella* genera (Smits et al., 2017). All isolations were performed under anaerobic conditions on YCFA-Glucose and YCFA-Baobab agar. Visible colonies from the initial plates were identified via colony PCR and re-plated onto BBE and LKV plates (Anaerobe Systems).

### Whole Genome Sequencing

Genomic DNA was extracted from single-isolate cultures grown for 24h using a MasterPure Gram Positive DNA Purification Kit. Long-read sequencing was performed using a Nanopore MinION and short read sequencing was performed using an Illumina MiSeq. Short read sequence quality was assessed using Fastqc with the command “fastqc --nogroup -q”, and adapters were trimmed with BBTools using the command “bbduk.sh -Xmx2g -eoom ref=adapters, phix threads=8 ktrim=r k=23 mink=11 edist=2 entropy=0.05 tpe tbo qtrim=rl minlength=100 trimq=30 pigz=t unpigz=t samplerate=0.25.” If there was more than 100x coverage of the genome, reads were normalized using the command “bbnorm.sh target=100 min=2”. Hybrid assembly of the short and long reads was performed using SPAdes with the command “spades.py —careful --cov-cutoff auto -k 21,33,55,77,99,127” (Prjibelski et al., 2020).

RagOUT was used for chromosome-level scaffolding using either the matched reference genome of the same species for *Bacteroides* (Table 1), or *Pc H-2477* for *Prevotella* (Kolmogorov et al., 2018).

Assembly quality was assessed with Quast (Mikheenko et al., 2018). Gene annotation was performed using RASTtk (Brettin et al., 2015).

### Clustering Genomes into Subspecies

All public *Bacteroides* and *Prevotella* genomes were downloaded from NCBI GenBank on 1/26/2021 using the program ncbi-genome-download (https://github.com/kblin/ncbi-genome-download). For *Bacteroides*, all genomes marked as “representative genome” in RefSeq (n=53) and genomes marked as assembly level “Complete Genome” or “Chromosome” in Genbank (n=71) were retained for further analysis (n=113 genomes retained of 1,229 total genomes). For *Prevotella*, all available public genomes were retained (n=368). Public genomes were clustered along with the isolate genomes recovered in the study using dRep v3.2.1(Olm et al., 2017) using the command “dRep dereplicate –S_algorithm fastANI - sa 0.98 –SkipMash -nc 0.65” to ensure that genomes with ≥ 98% ANI and ≥ 65% alignment coverage according to FastANI (Jain et al., 2018) are considered to be the same “subspecies”. These specific thresholds were chosen manually based on histograms of reported ANI and alignment coverage values. Representative genomes were chosen using dRep’s default scoring system with the following adjustments: public genomes marked as “representative genome” in Refseq were given an additional 50 points, and genomes recovered in this study were given an additional 200 points.

### Evaluating Subspecies Prevalence and Phylogenetic Analysis

Metagenomic reads were downloaded from Merrill et. al. (Merrill et al., 2022) (all other populations). Metagenomic reads were mapped to *Prevotella* and *Bacteroides* subspecies representative genomes using Bowtie2 (Langmead and Salzberg, 2012), and the resulting .bam files were profiled using inStrain (Olm et al., 2021). Genomes detected with ≥ 65% genome breadth were considered “present” in a metagenome. The prevalence of each genome in each population was calculated as the percentage of metagenomes in which the genome was detected.Phylogenetic trees were made all for *Bacteroides* and *Prevotella* subspecies representative genomes detected in at least one metagenome using GToTree v1.5.36 with the command “GToTree -H Bacteria”. One outgroup from a different genus was included in each tree. Tree leaves were labeled based on GTDB taxonomy release 202 (Chaumeil et al., 2020), which in some cases classified genomes as belonging to other genera than they were deposited in in GenBank. Trees were visualized using iTol (Letunic and Bork, 2021).

### CAZyme Annotation

CAZyme annotations were performed for each isolate. An additional 20 strains of *Prevotella copri* available at NCBI, with variable assembly levels, were annotated as well for comparative purpose, with the isolates and two model strains. All amino acid sequences were first compared to the full-length sequences stored in the CAZy database (Sept. 2021) (Drula et al., 2022) using BlastP (version 2.3.0+) (Camacho et al., 2009). Queries obtaining 100% coverage, >50% sequence identity and E-value ≤10^−6^ were automatically annotated with the same domain composition as the closest reference homolog. All remaining sequences were subject to human curation to verify the presence of each putative modules. During this process, the curator could rely on (i) bioinformatics tools, including BLAST against libraries on either full-length protein, modules only or characterized modules only, and HMMER version 3.1 (Mistry et al., 2013) against in-house built models for each CAZy (sub)family; (ii) human expertise on the appropriate coverage, sequence identity and E-value thresholds which vary across (sub)families, and ultimately on the verification of the catalytic amino-acid conservation. Hierarchical clustering of isolates’ CAZyme repertoires was performed using ComplexHeatmap (Gu, 2022). Predicted substrate assignment was compiled from previously published works (Desai et al., 2016; Smits et al., 2017).

### *In Vitro* Polysaccharide Growth Assays

Glycerol stocks were struck out on Brain Heart Infusion plates with 10% defibrinated horse blood and incubated anaerobically for 24 h at 37ºC. Isolates were passaged overnight in BHI-S (*Bacteroides*), and YCFA-G (*Prevotella*). After 16h, cultures were diluted 1:50 for *Bacteroides* and 1:10 for *Prevotella* into 200uL of culture media in a clear, flat bottomed 96-well plate. Growth media was composed of a YCFA background, plus 0.5% carbohydrate, with the exception of inulin, which was added at a 1.5% concentration. OD600 was measured every 15 minutes for 48h using a BioTek Epoch2 plate reader, with 30 seconds of shaking prior to each reading. Normalized OD was calculated for each carbohydrate condition by subtracting the average blank OD600 from the raw OD600 for each isolate grown in the corresponding polysaccharide. Maximum OD was calculated as the highest normalized OD in the first 24h period.

### Colonization and Enumeration of Gnotobiotic Mice

For colonization with *B. thetaiotaomicron H-2622*, mice were gavaged with 300uL of a 3mL culture grown for 16h in BHI-S. For colonization with *P. copri*, mice were gavaged with 300uL of a 3mL culture grown for 16h in YCFAC, in which was suspended 10-15 lawns (∼1 per mouse) of *P. copri* grown on BHIBA for 48 hours. For *Prevotella* colonization, food removed from mouse cages and bedding changed 12h before gavage. Before the gavage of *Prevotella*, mice were gavaged with 300uL of 10% sodium bicarbonate in water. Food was returned 2h post-gavage.

For bicolonization experiments, mice were first colonized with *Pc H-2477*, then gavaged with *Bt H-2622* 7 days later. Bicolonization was allowed to stabilize for 5-7 days before the diet switch. Feces were collected from individual mice. Two biological replicates of 1 μl feces were resuspended in 200 μl sterile PBS, serially diluted 1:10, and 2μl of each dilution was plated on BHIBA. CFUs were counted after 36h anaerobic growth at 37 □ °C.

### *In Vivo* Competition Assays

Feces were collected from individual mice. Genomic DNA was extracted from 2 biological replicates of fecal pellets using DNeasy PowerLyzer PowerSoil kit (Qiagen). Concentration of *Pc* and *Bt* DNA was assessed using species-specific qPCR primers (Key Resources Table). qPCR was performed using the Brilliant III, Ultra Fast SYBR Green QPCR Master Mix and a Bio Rad CFX thermocycler. Genomic DNA from *Bt H-2622* and *Pc H-2477* were used to generate a standard curve for each primer pair. The standard curves were used to calculate the absolute quantity of *Bt* or *Pc* DNA in the sample. The efficiency value (E) for each primer pair was calculated as 10^(1/−slope)^ of log10(DNA input) against Ct value. Competitive index was calculated using this equation: 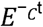 *Pc* primer pair/ 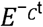 *Bt* primer pair. A competitive index of 1 denotes equal abundance.

### Mouse Diets

The Inulin-g and No MAC-g diets were created using 32% AIN-93G Basal Mix (CHO, Cellulose Free) and 68% carbohydrates, to match the carbohydrate content of the No MAC diet (TD.150689). The Basal Mix and carbohydrate components were suspended in a mixture of water (1100ml per 250g package of Basal Mix) and 5% bovine gelatin as a binder. The carbohydrates (100% glucose, no MAC-g; 50% glucose and 50% inulin, Inulin-g) and gelatin were dissolved separately in MilliQ water and autoclaved. The gelatin mix and AIN-93G Basal Mix (CHO, Cellulose Free) (TD.200788) were added to the carbohydrate solution in a tissue culture hood, and the mix was allowed to solidify at 4ºC. Diets are listed in the Key Resources Table. After one week post-colonization, standard chow was removed and replaced with the desired test diet, and the bedding was changed. Gelatin chow was replaced every 3 days as the chow dried out.

### RNAseq

RNA was extracted from mouse cecal contents and *in vitro* cultures using the RNeasy PowerMicrobiome Kit (Qiagen). Ribosomal RNA depletion was performed using the RiboMinus™ Transcriptome Isolation Kit (Invitrogen). A cDNA library was constructed using the TruSeq® Stranded Total RNA Library Prep Human/Mouse/Rat kit. Sequencing was performed on a NovaSeq SP flow cell.

Quality of raw reads was assessed with Multiqc using the command “multiqc”(Ewels et al., 2016). Adapters were trimmed using Trimmomatic and the command “trimmomatic PE ILLUMINACLIP - PE.fa:2:30:10 LEADING:3 TRAILING:3 SLIDINGWINDOW:4:15 MINLEN:36” (Bolger et al., 2014). Reads were aligned to the *Pc H-2477* and *Bt H-2622* genomes using HiSAT2 commands “hisat2-build” to generate indexes, and “hisat2 -p 8 --dta -x” to align reads to the indexes (Kim et al., 2019). SAMtools was used to generate .bam files with the commands “ samtools sort -@ 8 -o” and “samtools index” (Danecek et al., 2021). Transcripts were assembled using the Stringtie commands “stringtie”, “stringtie-merge”, and “stringtie -e -B -p 11 -G” (Pertea et al., 2016). Differential expression was analyzed using DESeq2 (Love et al., 2014).

---

**Figure S1.**
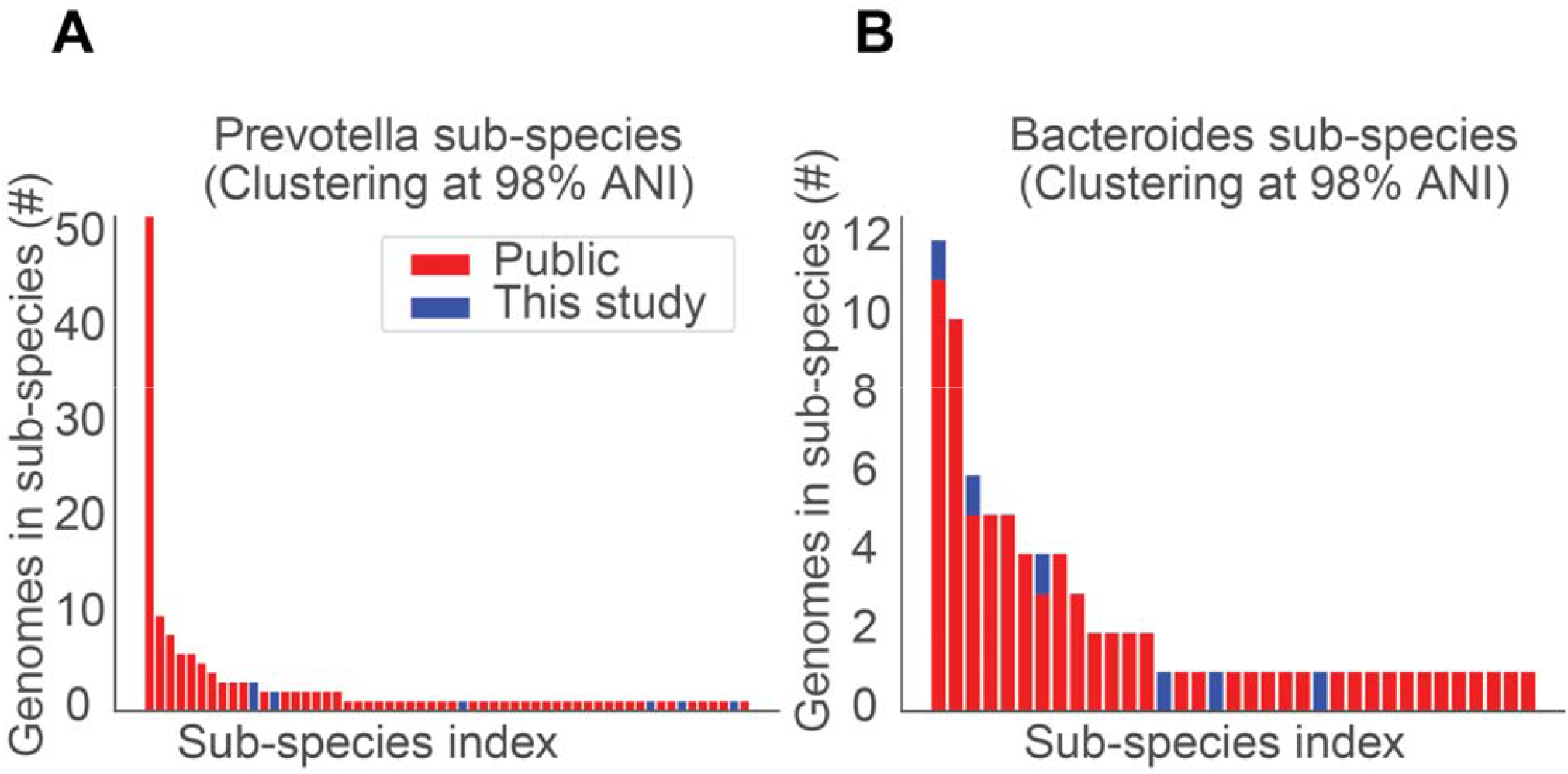
Clustering subspecies of *Bacteroides* and *Prevotella*. Number of representative genomes from *Prevotella* **(A)** and *Bacteroides* **(B)** subspecies used in the genome comparisons in **Fig. 1**. Sub-species index: genomes were clustered at 98% average nucleotide identity (ANI). Each bar represents one of these subspecies bins. Blue indicates genomes isolated for this study.

**Figure S2.**
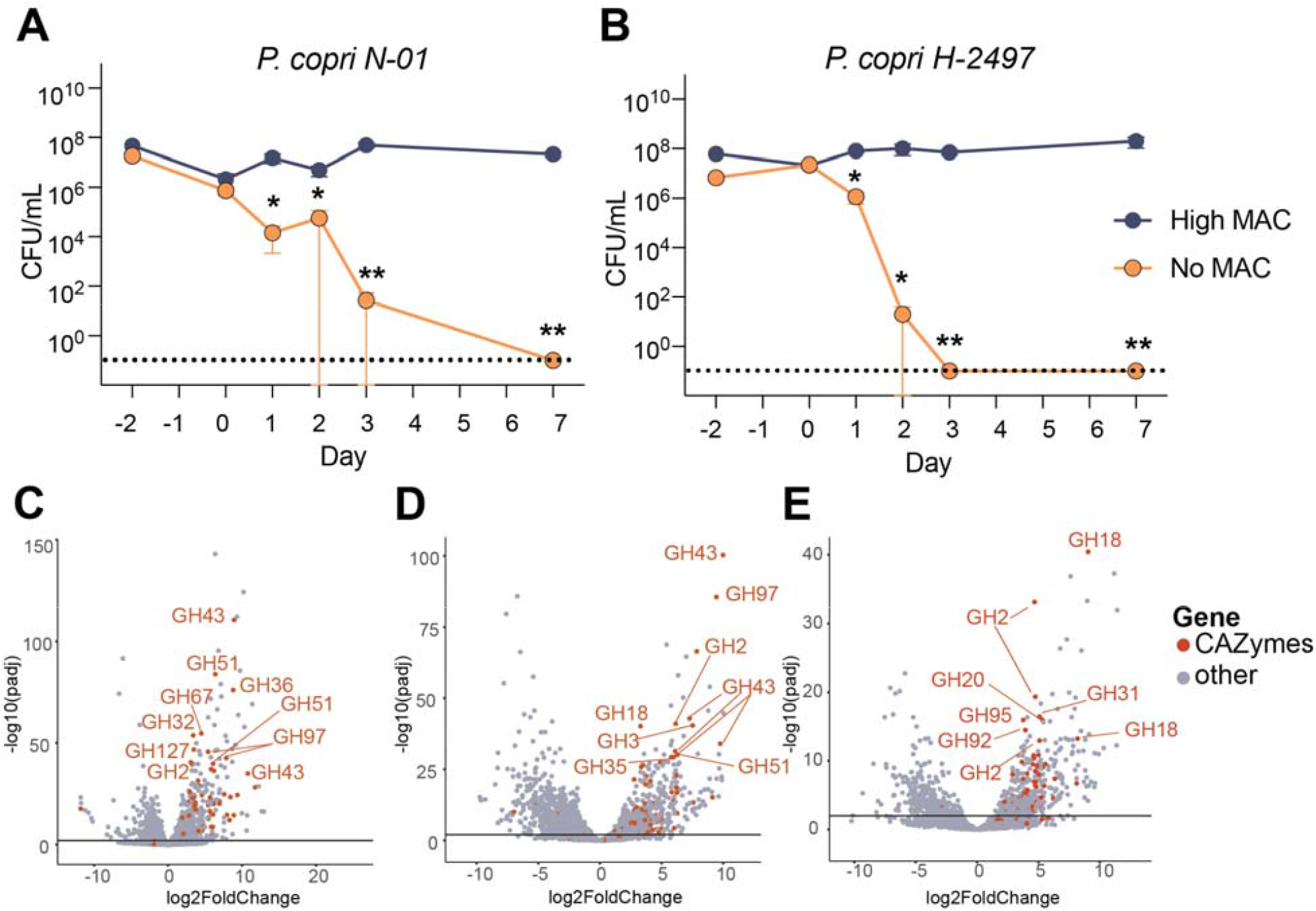
Diet-driven changes in *Bacteroides* and *Prevotella* colonization and gene expression. **(A, B)** Fecal density of *Pc N-01* **(A)**, and Hadza *Pc H-2497* **(B)**, in monocolonized mice fed different diets (Multiple Mann-Whitney tests, *: p ≤ 0.05, **: p ≤ 0.01). Dotted line indicates 0 CFU. Diet change to no MAC occurred on day 0. (**C, D)** Differential expression of *Pc H-2477* **(C)** or *Bt H-2622* **(D)** genes on High MAC *in vivo* vs. PYG *in vitro*. Black line at y=2. CAZyme genes marked in red. **(E)** Differential expression of *Bt H-2622* genes on No MAC *in vivo* vs. PYG *in vitro*. Black line at y=2. CAZyme genes marked in red.

**Figure S3.**
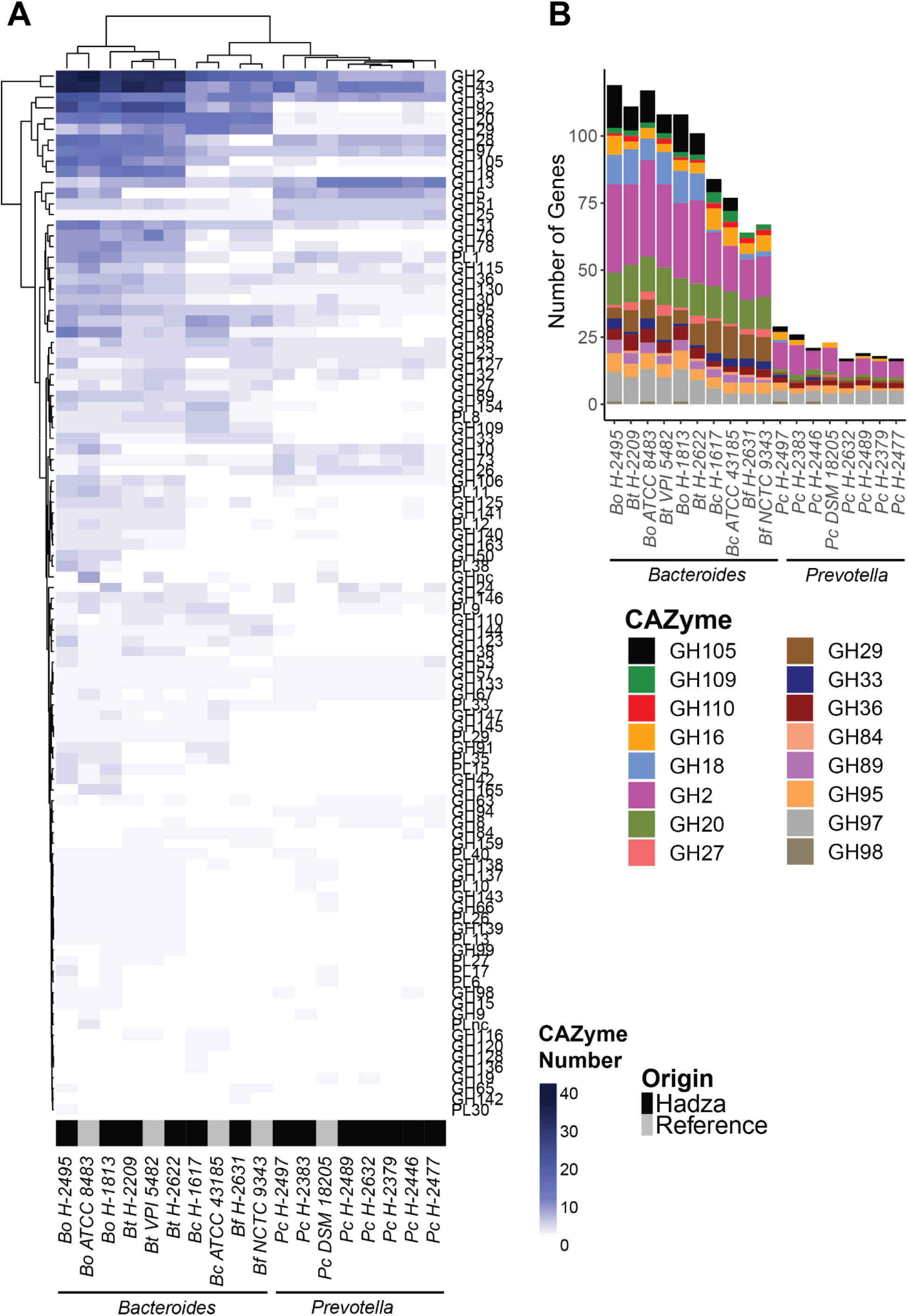
Extended CAZyme data for Hadza and reference isolates. **(A)** Number of all GHs and PLs per genome. Color gradient indicates the raw numbers of individual CAZymes of each family present in each genome. Hadza isolates indicated in black, reference strains indicated in gray. **(B)** Number of GHs from all putative mucin-degrading GH families.

## Notes

https://doi.org/10.5281/zenodo.7651179

## References

Abdill, R.J., Adamowicz, E.M., Blekhman, R., 2022. Public human microbiome data are dominated by highly developed countries. PLOS Biol. 20, e3001536. https://doi.org/10.1371/journal.pbio.3001536

Accetto, T., Avguštin, G., 2015. Polysaccharide utilization locus and CAZYme genome repertoires reveal diverse ecological adaptation of Prevotella species. Syst. Appl. Microbiol. 38, 453–461. https://doi.org/10.1016/J.SYAPM.2015.07.007

Bell, A., Juge, N., 2021. Mucosal glycan degradation of the host by the gut microbiota. Glycobiology 31, 691–696. https://doi.org/10.1093/GLYCOB/CWAA097 Bjursell, M.K., Martens, E.C., Gordon, J.I., 2006. Functional Genomic and Metabolic Studies of the Adaptations of a Prominent Adult Human Gut Symbiont, Bacteroides thetaiotaomicron, to the Suckling Period J. Biol. Chem. 281, 36269. https://doi.org/10.1074/jbc.M606509200

Bolger, A.M., Lohse, M., Usadel, B., 2014. Trimmomatic: A flexible trimmer for Illumina sequence data. Bioinformatics 30, 2114–2120. https://doi.org/10.1093/bioinformatics/btu170

Brettin, T., Davis, J.J., Disz, T., Edwards, R.A., Gerdes, S., Olsen, G.J., Olson, R., Overbeek, R., Parrello, B., Pusch, G.D., Shukla, M., Thomason, J.A., Stevens, R., Vonstein, V., Wattam, A.R., Xia, F., 2015. RASTtk: A modular and extensible implementation of the RAST algorithm for building custom annotation pipelines and annotating batches of genomes. Sci. Rep. 5, 8365. https://doi.org/10.1038/srep08365

Camacho, C., Coulouris, G., Avagyan, V., Ma, N., Papadopoulos, J., Bealer, K., Madden, T.L., 2009. BLAST+: architecture and applications. BMC Bioinformatics 10, 421. https://doi.org/10.1186/1471-2105-10-421

Chaumeil, P.-A., Mussig, A.J., Hugenholtz, P., Parks, D.H., 2020. GTDB-Tk: a toolkit to classify genomes with the Genome Taxonomy Database. Bioinformatics 36, 1925–1927. https://doi.org/10.1093/bioinformatics/btz848

Cordain, L., Eaton, S.B., Sebastian, A., Mann, N., Lindeberg, S., Watkins, B.A., O’Keefe, J.H., Brand-Miller, J., 2005. Origins and evolution of the Western diet: health implications for the 21st century. Am. J. Clin. Nutr. 81, 341–354. https://doi.org/10.1093/ajcn.81.2.341

Danecek, P., Bonfield, J.K., Liddle, J., Marshall, J., Ohan, V., Pollard, M.O., Whitwham, A., Keane, T., McCarthy, S.A., Davies, R.M., Li, H., 2021. Twelve years of SAMtools and BCFtools. GigaScience 10, giab008. https://doi.org/10.1093/gigascience/giab008

De Filippis, F., Pasolli, E., Tett, A., Tarallo, S., Naccarati, A., De Angelis, M., Neviani, E., Cocolin, L., Gobbetti, M., Segata, N., Ercolini, D., 2019. Distinct Genetic and Functional Traits of Human Intestinal Prevotella copri Strains Are Associated with Different Habitual Diets. Cell Host Microbe 25, 444-453.e3. https://doi.org/10.1016/j.chom.2019.01.004

De Filippo, C., Cavalieri, D., Di Paola, M., Ramazzotti, M., Poullet, J.B., Massart, S., Collini, S., Pieraccini, G., Lionetti, P., 2010. Impact of diet in shaping gut microbiota revealed by a comparative study in children from Europe and rural Africa. Proc. Natl. Acad. Sci. U. S. A. 107, 14691–6. https://doi.org/10.1073/pnas.1005963107

Desai, M.S., Seekatz, A.M., Koropatkin, N.M., Kamada, N., Hickey, C.A., Wolter, M., Pudlo, N.A., Kitamoto, S., Terrapon, N., Muller, A., Young, V.B., Henrissat, B., Wilmes, P., Stappenbeck, T.S., Núñez, G., Martens, E.C., 2016. A Dietary Fiber-Deprived Gut Microbiota Degrades the Colonic Mucus Barrier and Enhances Pathogen Susceptibility. Cell 167, 1339-1353.e21. https://doi.org/10.1016/J.CELL.2016.10.043

Dodd, D., Moon, Y.-H., Swaminathan, K., Mackie, R.I., Cann, I.K.O., 2010. Transcriptomic analyses of xylan degradation by Prevotella bryantii and insights into energy acquisition by xylanolytic bacteroidetes. J. Biol. Chem. 285, 30261–73. https://doi.org/10.1074/jbc.M110.141788

Drula, E., Garron, M.-L., Dogan, S., Lombard, V., Henrissat, B., Terrapon, N., 2022. The carbohydrate-active enzyme database: functions and literature. Nucleic Acids Res. 50, D571–D577. https://doi.org/10.1093/nar/gkab1045

Dubos, R.J., Pierce, C., 1948. The effect of diet on experimental tuberculosis of mice. Am. Rev. Tuberc. Pulm. Dis. 57, 287–293.

Earle, K.A., Billings, G., Sigal, M., Lichtman, J.S., Hansson, G.C., Elias, J.E., Amieva, M.R., Huang, K.C., Sonnenburg, J.L., 2015. Quantitative Imaging of Gut Microbiota Spatial Organization. Cell Host Microbe 18, 478–488. https://doi.org/10.1016/j.chom.2015.09.002

Ewels, P., Magnusson, M., Lundin, S., Käller, M., 2016. MultiQC: summarize analysis results for multiple tools and samples in a single report. Bioinformatics 32, 3047–3048. https://doi.org/10.1093/bioinformatics/btw354

Fehlner-Peach, H., Magnabosco, C., Raghavan, V., Scher, J.U., Tett, A., Cox, L.M., Gottsegen, C., Watters, A., Wiltshire-Gordon, J.D., Segata, N., Bonneau, R., Littman, D.R., 2019. Distinct Polysaccharide Utilization Profiles of Human Intestinal Prevotella copri Isolates. Cell Host Microbe 26, 680-690.e5. https://doi.org/10.1016/j.chom.2019.10.013

Flint, H.J., Scott, K.P., Duncan, S.H., Louis, P., Forano, E., 2012. Microbial degradation of complex carbohydrates in the gut. https://doi.org/10.4161/gmic.19897 3, 289–306. https://doi.org/10.4161/GMIC.19897

Fragiadakis, G.K., Smits, S.A., Sonnenburg, E.D., Van Treuren, W., Reid, G., Knight, R., Manjurano, A., Changalucha, J., Dominguez-Bello, M.G., Leach, J., Sonnenburg, J.L., 2019. Links between environment, diet, and the hunter-gatherer microbiome. Gut Microbes 10, 216–227. https://doi.org/10.1080/19490976.2018.1494103

Gálvez, E.J.C., Iljazovic, A., Amend, L., Lesker, T.R., Renault, T., Thiemann, S., Hao, L., Roy, U., Gronow, A., Charpentier, E., Strowig, T., 2020. Distinct Polysaccharide Utilization Determines Interspecies Competition between Intestinal Prevotella spp. Cell Host Microbe 28, 838-852.e6. https://doi.org/10.1016/j.chom.2020.09.012

Gorvitovskaia, A., Holmes, S.P., Huse, S.M., 2016. Interpreting Prevotella and Bacteroides as biomarkers of diet and lifestyle. Microbiome 4, 15. https://doi.org/10.1186/s40168-016-0160-7

Green, E.D., Gunter, C., Biesecker, L.G., Di Francesco, V., Easter, C.L., Feingold, E.A., Felsenfeld, A.L., Kaufman, D.J., Ostrander, E.A., Pavan, W.J., Phillippy, A.M., Wise, A.L., Dayal, J.G., Kish, B.J., Mandich, A., Wellington, C.R., Wetterstrand, K.A., Bates, S.A., Leja, D., Vasquez, S., Gahl, W.A., Graham, B.J., Kastner, D.L., Liu, P., Rodriguez, L.L., Solomon, B.D., Bonham, V.L., Brody, L.C., Hutter, C.M., Manolio, T.A., 2020. Strategic vision for improving human health at The Forefront of Genomics. Nature 586, 683–692. https://doi.org/10.1038/s41586-020-2817-4

Gu, Z., 2022. Complex heatmap visualization. iMeta 1, e43. https://doi.org/10.1002/imt2.43

Gu, Z., Gu, L., Eils, R., Schlesner, M., Brors, B., 2014. circlize implements and enhances circular visualization in R. Bioinformatics 30, 2811–2812. https://doi.org/10.1093/bioinformatics/btu393

Hahsler, M., Hornik, K., Buchta, C., 2008. Getting Things in Order: An Introduction to the R Package seriation. J. Stat. Softw. 25, 1–34. https://doi.org/10.18637/jss.v025.i03

Jain, C., Rodriguez-R, L.M., Phillippy, A.M., Konstantinidis, K.T., Aluru, S., 2018. High throughput ANI analysis of 90K prokaryotic genomes reveals clear species boundaries. Nat. Commun. 9, 5114. https://doi.org/10.1038/s41467-018-07641-9

Jha, A.R., Davenport, E.R., Gautam, Y., Bhandari, D., Tandukar, S., Ng, K.M., Fragiadakis, G.K., Holmes, S., Gautam, G.P., Leach, J., Sherchand, J.B., Bustamante, C.D., Sonnenburg, J.L., 2018. Gut microbiome transition across a lifestyle gradient in Himalaya. PLOS Biol. 16, e2005396. https://doi.org/10.1371/journal.pbio.2005396

Kaplan, R.C., Wang, Z., Usyk, M., Sotres-Alvarez, D., Daviglus, M.L., Schneiderman, N., Talavera, G.A., Gellman, M.D., Thyagarajan, B., Moon, J.-Y., Vázquez-Baeza, Y., McDonald, D., Williams-Nguyen, J.S., Wu, M.C., North, K.E., Shaffer, J., Sollecito, C.C., Qi, Q., Isasi, C.R., Wang, T., Knight, R., Burk, R.D., 2019. Gut microbiome composition in the Hispanic Community Health Study/Study of Latinos is shaped by geographic relocation, environmental factors, and obesity. Genome Biol. 20, 219. https://doi.org/10.1186/s13059-019-1831-z

Kim, D., Paggi, J.M., Park, C., Bennett, C., Salzberg, S.L., 2019. Graph-based genome alignment and genotyping with HISAT2 and HISAT-genotype. Nat. Biotechnol. 37, 907–915. https://doi.org/10.1038/s41587-019-0201-4

Kolmogorov, M., Armstrong, J., Raney, B.J., Streeter, I., Dunn, M., Yang, F., Odom, D., Flicek, P., Keane, T.M., Thybert, D., Paten, B., Pham, S., 2018. Chromosome assembly of large and complex genomes using multiple references. Genome Res. 28, 1720–1732. https://doi.org/10.1101/gr.236273.118

Langmead, B., Salzberg, S.L., 2012. Fast gapped-read alignment with Bowtie 2. Nat. Methods 9, 357–359. https://doi.org/10.1038/nmeth.1923

Lee, M.D., 2019. GToTree: a user-friendly workflow for phylogenomics. Bioinformatics 35, 4162–4164. https://doi.org/10.1093/bioinformatics/btz188

Letunic, I., Bork, P., 2021. Interactive Tree Of Life (iTOL) v5: an online tool for phylogenetic tree display and annotation. Nucleic Acids Res. 49, W293–W296. https://doi.org/10.1093/nar/gkab301

Li, J., Gálvez, E.J.C., Amend, L., Almási, É., Iljazovic, A., Lesker, T.R., Bielecka, A.A., Schorr, E.-M., Strowig, T., 2021. A versatile genetic toolbox for Prevotella copri enables studying polysaccharide utilization systems. EMBO J. 40, e108287. https://doi.org/10.15252/embj.2021108287

Love, M.I., Huber, W., Anders, S., 2014. Moderated estimation of fold change and dispersion for RNA-seq data with DESeq2. Genome Biol. 15, 550. https://doi.org/10.1186/s13059-014-0550-8

Luis, A.S., Jin, C., Pereira, G.V., Glowacki, R.W.P., Gugel, S.R., Singh, S., Byrne, D.P., Pudlo, N.A., London, J.A., Baslé, A., Reihill, M., Oscarson, S., Eyers, P.A., Czjzek, M., Michel, G., Barbeyron, T., Yates, E.A., Hansson, G.C., Karlsson, N.G., Cartmell, A., Martens, E.C., 2021. A single sulfatase is required to access colonic mucin by a gut bacterium. Nat. 2021 5987880 598, 332–337. https://doi.org/10.1038/s41586-021-03967-5

Marlowe, F.W., Berbesque, J.C., 2009. Tubers as fallback foods and their impact on Hadza hunter-gatherers. Am. J. Phys. Anthropol. 140, 751–758. https://doi.org/10.1002/ajpa.21040

Martens, E.C., Neumann, M., Desai, M.S., 2018. Interactions of commensal and pathogenic microorganisms with the intestinal mucosal barrier. Nat. Rev. Microbiol. 16, 457–470. https://doi.org/10.1038/s41579-018-0036-x

Merrill, B.D., Carter, M.M., Olm, M.R., Dahan, D., Tripathi, S., Spencer, S.P., Yu, B., Jain, S., Neff, N., Jha, A.R., Sonnenburg, E.D., Sonnenburg, J.L., 2022. Ultra-deep Sequencing of Hadza Hunter-Gatherers Recovers Vanishing Gut Microbes. https://doi.org/10.1101/2022.03.30.486478

Mikheenko, A., Prjibelski, A., Saveliev, V., Antipov, D., Gurevich, A., 2018. Versatile genome assembly evaluation with QUAST-LG. Bioinformatics 34, i142–i150. https://doi.org/10.1093/bioinformatics/bty266

Mistry, J., Finn, R.D., Eddy, S.R., Bateman, A., Punta, M., 2013. Challenges in homology search: HMMER3 and convergent evolution of coiled-coil regions. Nucleic Acids Res. 41, e121. https://doi.org/10.1093/nar/gkt263

Monteiro, C.A., Moubarac, J.-C., Cannon, G., Ng, S.W., Popkin, B., 2013. Ultra-processed products are becoming dominant in the global food system. Obes. Rev. 14, 21–28. https://doi.org/10.1111/obr.12107

Olm, M.R., Brown, C.T., Brooks, B., Banfield, J.F., 2017. dRep: a tool for fast and accurate genomic comparisons that enables improved genome recovery from metagenomes through de-replication. ISME J. 11, 2864–2868. https://doi.org/10.1038/ismej.2017.126

Olm, M.R., Crits-Christoph, A., Bouma-Gregson, K., Firek, B.A., Morowitz, M.J., Banfield, J.F., 2021. inStrain profiles population microdiversity from metagenomic data and sensitively detects shared microbial strains. Nat. Biotechnol. 39, 727–736. https://doi.org/10.1038/s41587-020-00797-0

Olm, M.R., Dahan, D., Carter, M.M., Merrill, B.D., Yu, F.B., Jain, S., Meng, X., Tripathi, S., Wastyk, H., Neff, N., Holmes, S., Sonnenburg, E.D., Jha, A.R., Sonnenburg, J.L., 2022. Robust variation in infant gut microbiome assembly across a spectrum of lifestyles. Science 376, 1220–1223. https://doi.org/10.1126/science.abj2972

Pertea, M., Kim, D., Pertea, G.M., Leek, J.T., Salzberg, S.L., 2016. Transcript-level expression analysis of RNA-seq experiments with HISAT, StringTie and Ballgown. Nat. Protoc. 11, 1650–1667. https://doi.org/10.1038/nprot.2016.095

Prjibelski, A., Antipov, D., Meleshko, D., Lapidus, A., Korobeynikov, A., 2020. Using SPAdes De Novo Assembler. Curr. Protoc. Bioinforma. 70, e102. https://doi.org/10.1002/cpbi.102

Pruss, K.M., Sonnenburg, J.L., 2021. C. difficile exploits a host metabolite produced during toxin-mediated disease. Nat. 2021 5937858 593, 261–265. https://doi.org/10.1038/s41586-021-03502-6

Pudlo, N.A., Urs, K., Crawford, R., Pirani, A., Atherly, T., Jimenez, R., Terrapon, N., Henrissat, B., Peterson, D., Ziemer, C., Snitkin, E., Martens, E.C., 2022. Phenotypic and Genomic Diversification in Complex Carbohydrate-Degrading Human Gut Bacteria. mSystems 7. https://doi.org/10.1128/MSYSTEMS.00947-21

Sakai, R., Winand, R., Verbeiren, T., Moere, A.V., Aerts, J., 2014. dendsort: modular leaf ordering methods for dendrogram representations in R. https://doi.org/10.12688/f1000research.4784.1

Salyers, A.A., West, S.E., Vercellotti, J.R., Wilkins, T.D., 1977. Fermentation of mucins and plant polysaccharides by anaerobic bacteria from the human colon. Appl. Environ. Microbiol. 34, 529–533. https://doi.org/10.1128/aem.34.5.529-533.1977

Smits, S.A., Leach, J., Sonnenburg, E.D., Gonzalez, C.G., Lichtman, J.S., Reid, G., Knight, R., Manjurano, A., Changalucha, J., Elias, J.E., Dominguez-Bello, M.G., Sonnenburg, J.L., 2017. Seasonal Cycling in the Gut Microbiome of the Hadza Hunter-Gatherers of Tanzania Authors: Science 806, 1–18. https://doi.org/10.1126/science.aan4834

Sonnenburg, E.D., Smits, S.A., Tikhonov, M., Higginbottom, S.K., Wingreen, N.S., Sonnenburg, J.L., 2016. Diet-induced extinctions in the gut microbiota compound over generations. Nature 529, 212–215. https://doi.org/10.1038/nature16504

Sonnenburg, E.D., Sonnenburg, J.L., 2014. Starving our microbial self: The deleterious consequences of a diet deficient in microbiota-accessible carbohydrates. Cell Metab. 20, 779–786. https://doi.org/10.1016/j.cmet.2014.07.003

Sonnenburg, E.D., Zheng, H., Joglekar, P., Higginbottom, S.K., Firbank, S.J., Bolam, D.N., Sonnenburg, J.L., 2010. Specificity of Polysaccharide Use in Intestinal Bacteroides Species Determines Diet-Induced Microbiota Alterations. Cell 141, 1241–1252. https://doi.org/10.1016/j.cell.2010.05.005

Sonnenburg, J.L., Sonnenburg, E.D., 2019. Vulnerability of the industrialized microbiota. Science 366, eaaw9255. https://doi.org/10.1126/science.aaw9255

Sonnenburg, J.L., Xu, J., Leip, D.D., Chen, C.H., Westover, B.P., Weatherford, J., Buhler, J.D., Gordon, J.I., 2005. Glycan foraging in vivo by an intestine-adapted bacterial symbiont. Science 307, 1955\uc0\u8211{}1959. https://doi.org/10.1126/SCIENCE.1109051

Sprockett, D.D., Martin, M., Costello, E.K., Burns, A.R., Holmes, S.P., Gurven, M.D., Relman, D.A., 2020. Microbiota assembly, structure, and dynamics among Tsimane horticulturalists of the Bolivian Amazon. Nat. Commun. 11, 3772. https://doi.org/10.1038/s41467-020-17541-6

Tett, A., Huang, K.D., Asnicar, F., Fehlner-Peach, H., Pasolli, E., Karcher, N., Armanini, F., Manghi, P., Bonham, K., Zolfo, M., De Filippis, F., Magnabosco, C., Bonneau, R., Lusingu, J., Amuasi, J., Reinhard, K., Rattei, T., Boulund, F., Engstrand, L., Zink, A., Collado, M.C., Littman, D.R., Eibach, D., Ercolini, D., Rota-Stabelli, O., Huttenhower, C., Maixner, F., Segata, N., 2019. The Prevotella copri Complex Comprises Four Distinct Clades Underrepresented in Westernized Populations. Cell Host Microbe 0. https://doi.org/10.1016/j.chom.2019.08.018

Vangay, P., Johnson, A.J., Ward, T.L., Kashyap, P.C., Culhane-Pera, K.A., Knights Correspondence, D., 2018. US Immigration Westernizes the Human Gut Microbiome. https://doi.org/10.1016/j.cell.2018.10.029

Wibowo, M.C., Yang, Z., Borry, M., Hübner, A., Huang, K.D., Tierney, B.T., Zimmerman, S., Barajas-Olmos, F., Contreras-Cubas, C., García-Ortiz, H., Martínez-Hernández, A., Luber, J.M., Kirstahler, P., Blohm, T., Smiley, F.E., Arnold, R., Ballal, S.A., Pamp, S.J., Russ, J., Maixner, F., Rota-Stabelli, O., Segata, N., Reinhard, K., Orozco, L., Warinner, C., Snow, M., LeBlanc, S., Kostic, A.D., 2021. Reconstruction of ancient microbial genomes from the human gut. Nature 594, 234–239. https://doi.org/10.1038/s41586-021-03532-0

Wickham, H., 2016. Programming with ggplot2, in: Wickham, H. (Ed.), Ggplot2: Elegant Graphics for Data Analysis, Use R! Springer International Publishing, Cham, pp. 241–253. https://doi.org/10.1007/978-3-319-24277-4_12

Wickham, H., n.d. An SVG Graphics Device [WWW Document]. URL https://svglite.r-lib.org/ x(accessed 2.8.23).

Wickham, H., Averick, M., Bryan, J., Chang, W., McGowan, L.D., François, R., Grolemund, G., Hayes, A., Henry, L., Hester, J., Kuhn, M., Pedersen, T.L., Miller, E., Bache, S.M., Müller, K., Ooms, J., Robinson, D., Seidel, D.P., Spinu, V., Takahashi, K., Vaughan, D., Wilke, C., Woo, K., Yutani, H., 2019. Welcome to the Tidyverse. J. Open Source Softw. 4, 1686. https://doi.org/10.21105/joss.01686

Xu, J., Bjursell, M.K., Himrod, J., Deng, S., Carmichael, L.K., Chiang, H.C., Hooper, L.V., Gordon, J.I., 2003. A Genomic View of the Human-Bacteroides thetaiotaomicron Symbiosis. Science 299, 2074–2076. https://doi.org/10.1126/science.1080029

Yatsunenko, T., Rey, F.E., Manary, M.J., Trehan, I., Dominguez-Bello, M.G., Contreras, M., Magris, M., Hidalgo, G., Baldassano, R.N., Anokhin, A.P., Heath, A.C., Warner, B., Reeder, J., Kuczynski, J., Caporaso, J.G., Lozupone, C.A., Lauber, C., Clemente, J.C., Knights, D., Knight, R., Gordon, J.I., 2012. Human gut microbiome viewed across age and geography. Nature 486, 222–227. https://doi.org/10.1038/nature11053

